# CXCR6 promotes dermal CD8^+^ T cell survival and transition to long-term tissue residence

**DOI:** 10.1101/2023.02.14.528487

**Authors:** Taylor A. Heim, Ochapa Ibrahim, Ziyan Lin, Austin C. Schultz, Maria M. Steele, Tenny Mudianto, Amanda W. Lund

**Affiliations:** Ronald O. Perelman Department of Dermatology, NYU Grossman School of Medicine, New York, NY, USA; Applied Bioinformatics Laboratories, NYU Langone Health, New York, NY, USA; Department of Pathology, NYU Grossman School of Medicine, New York, NY, USA; Laura and Isaac Perlmutter Cancer Center, NYU Grossman School of Medicine, New York, NY, USA

## Abstract

Tissue resident memory T cells (T_RM_) provide protection against local re-infection, and yet the interstitial signals necessary for their formation and persistence remain incompletely understood. Here we show that antigen-dependent induction of the chemokine receptor, CXCR6, is a conserved adaptation to peripheral tissue infiltration that promotes T_RM_ formation after viral infection. Deficient T_RM_ formation in the absence of CXCR6 was not explained by canonical trafficking as CXCR6 was not required for tissue entry, was dispensable for the early accumulation of antigen-specific CD8^+^ T cells in skin, and did not restrain their exit. Further, single cell sequencing indicated that *Cxcr6*^-/-^ CD8^+^ T cells were competent to acquire a transcriptional program of residence and T_RM_ that formed were equally functional compared to their WT counterparts when reactivated greater than 100 days post primary infection. The reduced numbers observed at memory time points, where instead found to associate with impaired redox homeostasis and antioxidant capacity during the transition from effector to memory states. As such, *Cxcr6*^-/-^ CD8^+^ T cells exhibited increased rates of apoptosis in the dermis relative to controls, which led to reduced numbers of T_RM_ in the epidermis at memory. CXCR6 therefore promotes the metabolic adaptation of T cells as they engage antigen in tissue to increase the probability of memory differentiation and long-term residence.

**One Sentence Summary:** CXCR6 promotes mechanisms of cellular adaptation to tissue that support local survival and the transition to tissue residence.

## INTRODUCTION

Resident memory T cells (T_RM_) provide rapid, localized protection due to their position in barrier tissues. T_RM_ formation depends first upon the recruitment of effector T cells to sites of inflammation where local cues then direct the acquisition of a transcriptional program required for tissue adaptation and persistence. One key feature of this program is the downregulation of a transcriptional module associated with tissue exit via the lymphatic vasculature, including *Klf2*, *Ccr7*, and *S1pr1*, which presumably helps fix T_RM_ in a sedentary state. Consistent with this idea, *Ccr7* deficiency prevents CD4^+^ T cell egress from the skin via dermal lymphatic vessels following viral infection (1) and boosts CD8^+^ T_RM_ formation (2). Similarly, constitutive expression of *S1pr1* limits CD8^+^ T_RM_ formation in skin (3) and the conserved T_RM_ marker CD69, is thought to promote residency by sequestering S1PR1 and inhibiting T cell egress (4). While these mechanisms may limit the exit of T_RM_ progenitors from inflamed tissues, the signals that instruct T cell position within tissue along the T_RM_ differentiation trajectory, and the interstitial cues that switch between these migratory modules, remain incompletely understood.

Interestingly, one of the highly expressed and conserved chemokine receptors in both CD4^+^ and CD8^+^ T_RM_ across mice and humans is CXCR6 (5–7). The sole ligand for CXCR6, CXCL16 (8), is expressed by multiple cell types including macrophages, dendritic cells, keratinocytes, and endothelial cells (9–12). Induced by TNFα and IFNγ, CXCL16 is expressed in a transmembrane form and acts in trans at the cell-cell interface, but may also be cleaved from the cell surface to signal as a classical chemoattractant (13). CXCR6 loss impairs T_RM_ formation or maintenance in lung (14), skin (15), and liver (16), however, the specific mechanism by which CXCR6 guides CD8^+^ T_RM_ formation remains incompletely understood. In a mouse model of influenza infection, CXCR6 is dispensable for CD8^+^ T cell trafficking to the lung parenchyma but needed for localization to the CXCL16-expressing lung epithelium (14). In the liver, however, CXCR6 has been shown to be necessary for tissue infiltration during a graft vs host response (17), but also dispensable for early accumulation following malaria infection (16). Therefore, across peripheral, non-lymphoid tissues, when, exactly where, and how CXCR6 promotes CD8^+^ T cell accumulation remains unclear.

Here we used cutaneous, murine viral infection to define the kinetics of CXCR6 expression and its impact on the transition of effector CD8^+^ T cells to long-term tissue residence. We find that circulating CD8^+^ T cells in the spleen and within non-lymphoid peripheral tissues are distinguished from CD8^+^ T_RM_ in part by CXCR6 expression. CXCR6 expression was conserved across anatomical compartments where it was necessary for optimal CD8^+^ T_RM_ formation. In skin, rather than facilitating early CD8^+^ T cell accumulation or directly preventing egress, we demonstrate that local antigen encounter upregulates CXCR6 to promote CD8^+^ T cell survival by supporting redox homeostasis in the dermis allowing for more efficient seeding of the epidermis at memory. These data add to a mechanistic model for how CD8^+^ T cell position in tissue optimizes receipt of the signals necessary for T_RM_ formation and implicates CXCR6 as a conserved feature of tissue adaptation and survival.

## RESULTS

### CXCR6 optimizes CD8^+^ T_RM_ formation across peripheral, non-lymphoid tissues

In mice and humans *Cxcr6* is a part of the core cluster of genes associated with tissue residence (5–7), however, the mechanisms that regulate *Cxcr6* expression and its specific role in mediating peripheral tissue CD8^+^ T cell behavior remain unclear. To track CXCR6 expression and determine its functional contribution to T_RM_ formation we infected C57Bl/6 mice with vaccinia virus (VACV) by scarification on the ear pinna (10^6^ PFU). Infection with VACV by scarification peaks day 3-5 post infection, is cleared by day 15, and no viable virus is found at memory time points (**Fig S1A**) (18, 19). Consistent with this, there is no evidence that CD8^+^ T cells engage antigen in the skin 20 days after infection (20). As such, VACV infection by scarification is used to model the generation of anti-viral, long-lived skin tissue resident memory(19–21). Tracking anti-viral T cells in this model, we found that CXCR6 was highly expressed on the surface of endogenous VACV-specific CD8^+^ T cells (H2-K^b^-B8R_20-27_) in skin 30 days post infection (d.p.i). but was relatively low in VACV-specific and naive CD8^+^ T cells in the spleen at matched time points (**Fig 1A and B**). While CD103 labels a subset (38±7% of CD69^+^) of antigen-specific T_RM_ in skin 28 d.p.i., CXCR6 was highly expressed on a larger percentage of CD69^+^ cells (85±4%) (**Fig 1A**). Further, CD69^+^ cells, which are considered to be bona fide T_RM_ (19), had significantly higher levels of CXCR6 compared to CD69^-^ T cells in skin at the same time point (**Fig 1C**), all consistent with a role for CXCR6 in the formation of skin T_RM_. CXCR6 promotes the formation of T_RM_ in lung using an intramuscular vaccine with an adjuvant and antigen pull into the lung (14), supports accumulation of *ex vivo* activated T cells in skin in response to local dinitrofluorobenzene application (15), and promotes effector T cell function in tumors (22, 23). We therefore asked if CXCR6 promotes the *de novo* generation of T_RM_ post infection with VACV. To this end, we co-transferred equal numbers of congenically mismatched, WT and *Cxcr6*^-/-^ (*Cxcr6*^GFP/GFP^) (24) OT-1 T cells, specific for H2K^b^-OVA_257-264_ (SIINFEKL), into mice and infected ear skin with vaccinia expressing the immunodominant CD8^+^ T cell epitope of ovalbumin (OVA_257-264_; VACV-OVA) the following day (**Fig 1D and Fig S1B**). We compared the ratio of WT and CXCR6-/- OT-1 T cells more than 45 d.p.i. and observed significant enrichment for WT over *Cxcr6*^-/-^ OT-1 T cells in the skin relative to the spleen (**Fig 1E and F**). Interestingly, the memory advantage provided by CXCR6 was restricted to the resident compartment. Analysis of both VACV-specific CD69^-^ circulating memory and CD69^+^ T_RM_ at late memory time points (>100 d.p.i) revealed that only T_RM_ and not circulating memory, either in spleen or circulating through skin, were outcompeted by wildtype (**Fig 1G**). Interestingly, of the OT-1 T cells within the skin, WT cells exhibited more robust CD69 and CD103 expression than *Cxcr6*^-/-^ OT-1 T cells (**Fig S1C and D**), perhaps indicating a less differentiated T_RM_ phenotype. These data indicated that CXCR6 was required for optimal CD8^+^ T_RM_ formation in the skin following VACV infection.

**Figure 1.**
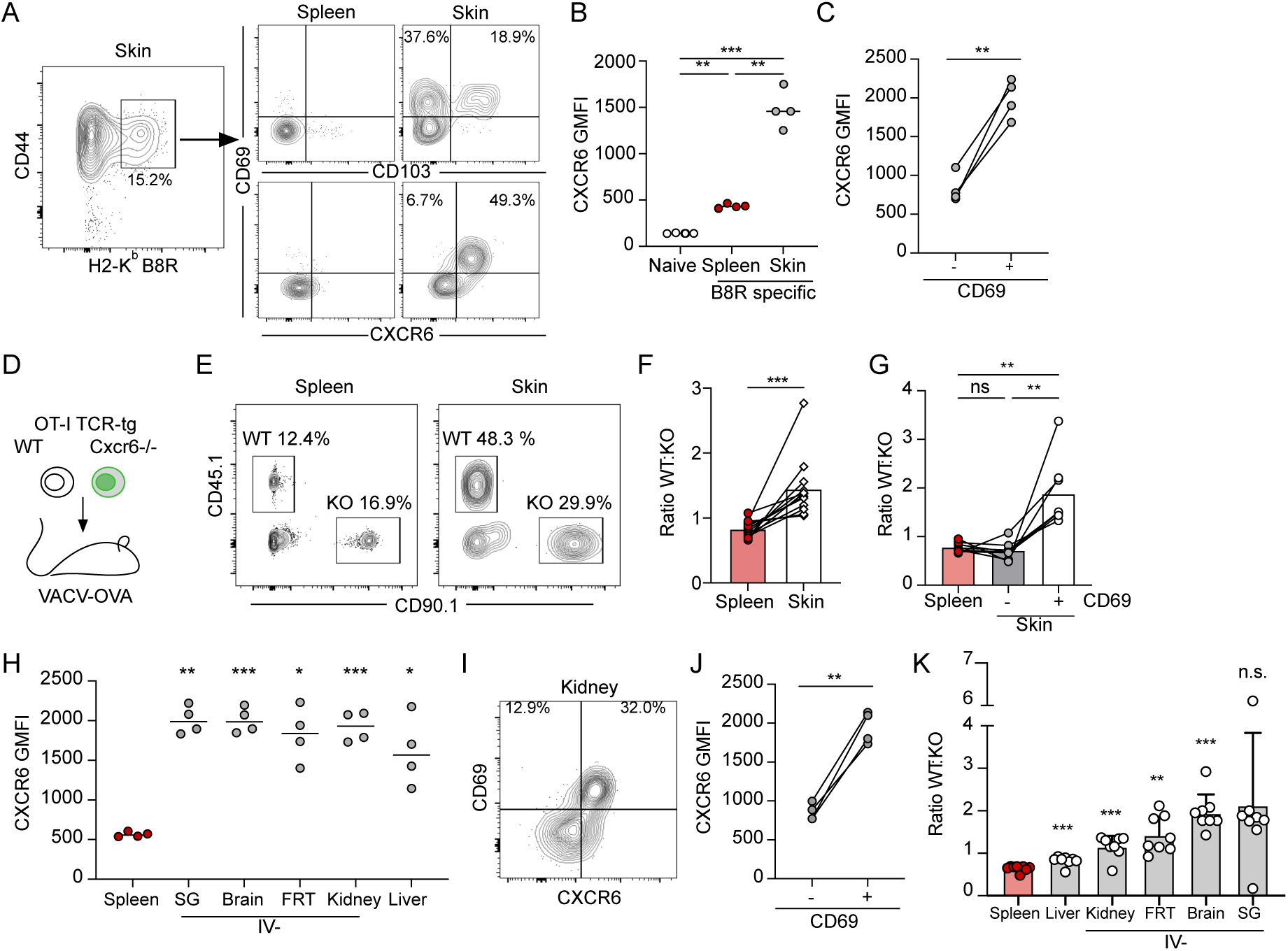
CXCR6 expression is a conserved feature of resident CD8^+^ T cells after viral infection. (**A**) Representative flow of CD44^+^ vaccinia virus (VACV) specific (H2-K^b^B8R_20-27_) CD8^+^ T cells 28 days post infection (d.p.i.) (VACV, scarification) and expression of CD69, CD103, and CXCR6. (**B, C**) Quantification of CXCR6 geometric mean fluorescence intensity (GMFI) on CD44^+^CD8^+^ B8R-specific T cells from (A), as a function of location (B) and CD69 expression (C). (**D**) Experimental design where 15,000 wildtype (WT) OT-1 T cells (CD45.1^+^CD90.1^-^) and 15,000 *Cxcr6*^-/-^ OT-1 T cells (CD45.1^-^CD90.1^+^) were co-transferred into naïve mice (CD45.1^-^ CD90.1^-^) and the following day, ear skin was infected with VACV expressing OVA_257-264_ (VACV-OVA, scarification). (**E**) Representative flow plots at least 49 days post infection (d.p.i.). Gated on CD8^+^CD44^+^IV^-^ in the skin. (**F**) Quantification of the ratio of WT and *Cxcr6*^-/-^ OT-1 T cells isolated from the spleen and ear skin from (**E**). (**E and F**) Data are cumulative from 3 experiments for a total of n=12. (**G**) Quantification of the ratio of WT and *Cxcr6*^-/-^ 100-115 days post VACV infection stratified by CD69 expression. (**H**) CXCR6 expression on CD45.1^+^ OT-1 T cells in non-lymphoid tissues 31 d.p.i. with vesicular stomatitis virus expressing OVA (VSV-OVA) (gated on CD8^+^CD45.1^+^CD44^+^CD69^+^, and IV^-^ in all tissues except liver and spleen) (SG = salivary gland and FRT = Female reproductive tract). (**I**) Representative flow plot and (**J**) quantification of CXCR6 GMFI on IV^-^CD45.1^+^ OT-1 T cells from kidney as a function of CD69 expression. (**K**) Quantification of the ratio of WT and *Cxcr6*^-/-^ OT-1 T cells isolated 31 d.p.i. with VSV-OVA. Gated on CD8^+^CD44^+^IV^-^ in kidney, FRT, brain and salivary gland; CD8^+^CD44^+^ in liver and spleen. Data are cumulative of two experiments for a total of n=8. Bars represent average + SD. (**A-G)** Data are representative of at least two experiments with n=4 each. Statistical significance determined with one-way ANOVA (**B, G, and H**), repeated measures one-way ANOVA (**K**), and paired t test (**C, F, and J**). * p < 0.05, ** p < 0.01, *** p < 0.001, **** p < 0.0001

Given that T_RM_ phenotype is highly dependent upon pathogenic insult and anatomical location (25), we asked whether high CXCR6 expression was restricted to CD8^+^ T_RM_ formed following cutaneous VACV infection using vesicular stomatitis virus (VSV), which establishes CD8^+^ T_RM_ in diverse non-lymphoid tissues when delivered intravenously (10^6^ PFU) (26). We first transferred naïve CD45.1^+^ OT-1 T cells into mice and infected them with VSV-OVA. As we saw in skin with VACV, in all tissues tested (liver, brain, kidney, salivary gland, and female reproductive tract), CXCR6 expression was higher on tissue CD8^+^ T_RM_ (at least 30 d.p.i.) than antigen-specific CD8^+^ T cells in the spleen (**Fig 1H**). Furthermore, CD69^+^ OT-1 T_RM_ expressed significantly more CXCR6 compared to CD69^-^ T cells in the same tissue (**Fig 1I and J**), suggesting that CXCR6 expression is a conserved feature of anti-viral, CD8^+^ T_RM_. Importantly, the accumulation of T_RM_ in other peripheral non-lymphoid tissues, including the brain, kidney and female reproductive tract, depended in part on CXCR6 expression (**Fig 1K**), with WT T cells outcompeting *Cxcr6*^-/-^ in all tissues examined, though differences in expression of CD69 and CD103 were tissue specific (**Fig S2E-I**). These data together indicate that CXCR6 optimizes *de novo* CD8^+^ T_RM_ formation in disparate anatomical locations and in response to diverse viral infections.

### CXCR6 expression is an early event along the T_RM_ differentiation trajectory

These data indicated that even with a fixed TCR and at a static time point, CXCR6 seemed to distinguish a pool of circulating and resident CD8^+^ T cells within the tissue interstitium. To gain higher resolution on the transcriptional CD8^+^ T cell states captured within tissue at a single time point and to understand where CXCR6 is turned on in the T_RM_ differentiation trajectory, we performed single-cell RNA sequencing (scRNAseq) on interstitial CD8^+^ T cells. Congenically distinct, OT-1 T cells were transferred intravenously into C57Bl/6 mice. The following day, mice were infected with VACV expressing OVA_257-264_ (VACV-OVA). Live, extravascular OT-1 T cells (negative for intravascular stain; IV^-^) were sorted from infected skin 21 d.p.i. and submitted for scRNAseq to capture cells at the beginning of memory formation (19). Transcriptional analysis and visualization using Uniform Manifold Approximation Projection (UMAP) identified six clusters (**Fig. 2A and Fig S2A and B**). Cluster 1 expressed high levels of *Tcf7, Klf2,* and *S1pr1* and scored for a core circulating gene signature (T_CIRC_, **Fig. S2C and D**), while clusters 2-5 expressed high levels of *Itgae* and *Cd69* and enriched for a transcriptional program associated with residence (T_RM_, **Fig. S2E and F**). Cluster 6, interestingly expressed high levels of *Zeb2*, a marker of terminal differentiation that is absent from long-lived T_RM_ memory following lymphocytic choriomeningitis virus (LCMV) infection (27). Cluster 6 also differentially expressed genes associated with RNA splicing and apoptotic cleavage of cellular proteins, perhaps indicating an increased rate in cell death (**Fig. S2B**). The heterogeneity observed within the T_RM_ clusters suggested that even at a single time point and with a fixed TCR, CD8^+^ T cells could be captured in distinct transcriptional states that might reflect either functional or positional heterogeneity within the tissue. We performed scVelo analysis to define a pseudotime trajectory between the clusters and this analysis indicated a directional differentiation path from the circulating cluster 1 towards the T_RM_ clusters progressing from the cluster 3 to 2 and 5 to 4 (**Fig. 2B-D**). Consistent with this trajectory, the T_RM_ subclusters (2–5) varied in their expression of effector molecule transcripts and several transcription factors related to T cell activation and differentiation (**Fig. S2A**). Cells in cluster 3, exhibited an intermediate state consistent with their position in pseudotime at the transition from circulation and residence. They did not express transcripts high enough to meet our criteria for marker genes (fold change of 2, relative to average expression in other clusters), but expressed the highest levels of *Klf2*, *Tcf7* and *S1pr1* of all of the T_RM_ clusters. Cluster 2 cells were defined by high expression of chemokines *Ccl3*, *Ccl4* and transcription factors *Fos* and *Jun*, which may be indicative of recent activation (28) and Cluster 5 cells expressed *Lmna* and *Anxa1*. The predicted terminal state, Cluster 4, was defined by highest expression of cytotoxic molecules including *Gzma* and *Gzmf,* and *Krt83*, a keratin primarily found in the hair follicle*. Cxcr6* expression was elevated in all T_RM_ subclusters relative to the circulating cluster 1, with high expression already in the intermediate cluster 3, pointing towards the induction of *Cxcr6* expression as an early event in T_RM_ differentiation (**Fig. 2E**). As previously reported, T_RM_ CD8^+^ T cells exhibited lower expression of *S1pr1* relative to T_CIRC_, consistent with its proposed role in tissue egress (3). Of note, little *Ccr7* expression was seen in any subset, instead T_CIRC_ cells expressed *Cxcr4,* which facilitates egress from melanoma through CXCL12-producing dermal lymphatic vessels (29). In addition to *Cxcr6*, several receptors including, *S1pr4*, *Ccr2*, *Ccr5,* and *Cxcr3* were enriched in T_RM_ relative to T_CIRC_ (**Fig S1G and H).** As all cells analyzed stained negative for an intravascular stain, these data all together begin to indicate that the high surface *Cxcr6* expression observed across T_RM_ likely depended upon a transcriptional event that occurred after tissue entry.

**Figure 2.**
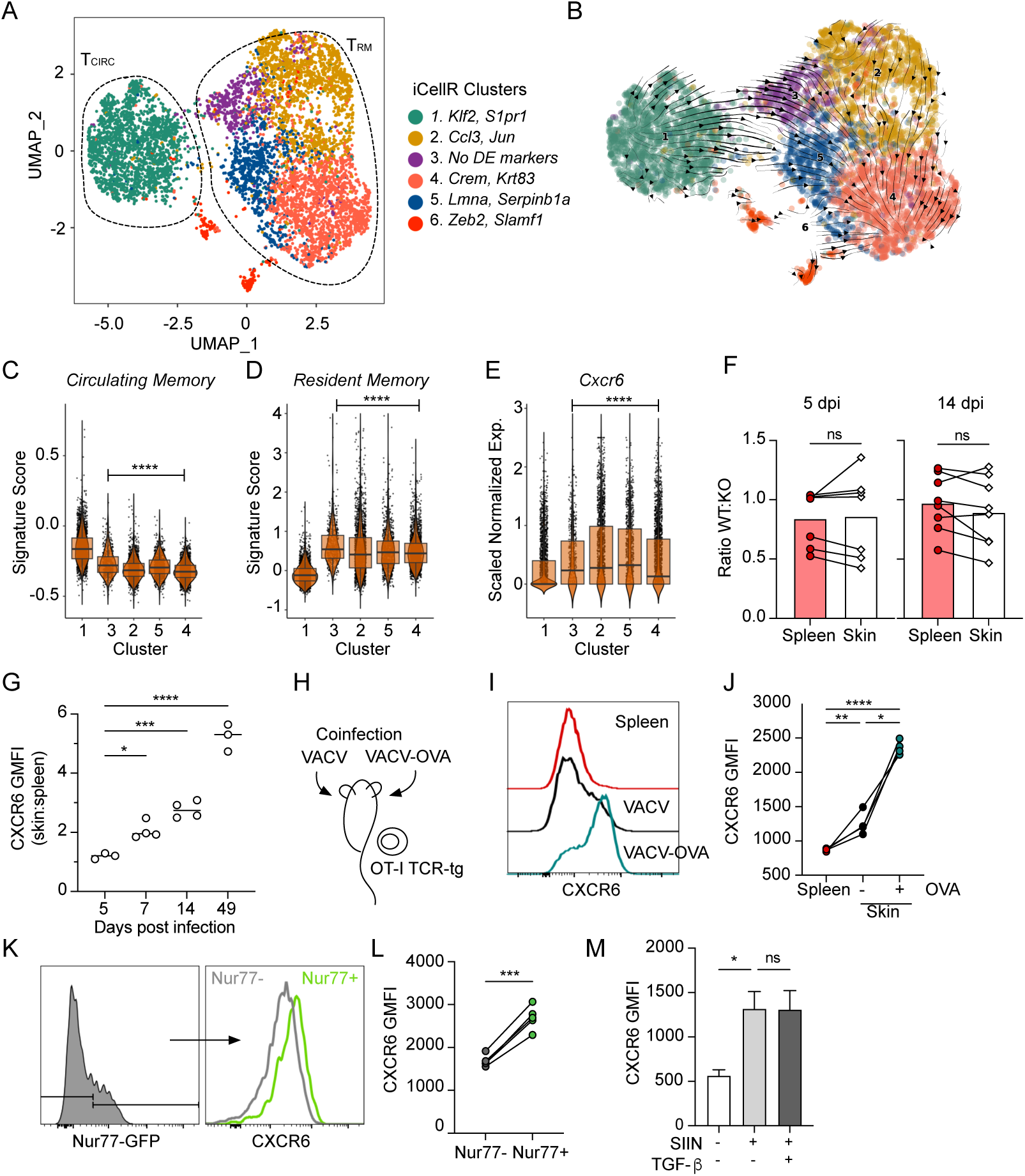
CXCR6 is required for optimal formation of T_RM_ across non-lymphoid peripheral tissues. (**A**) UMAP representation of single-cell RNA-seq data from extravascular (IV^-^) OT-1 CD8^+^ T cells isolated out of skin 21 days post infection (d.p.i.) with vaccinia virus expressing OVA_257-264_ (VACV-OVA). Circulating (T_circ_) and resident (T_RM_) memory T cells. Data represent a total of 7409 cells across all clusters. (**B**) Projected RNA velocity (scVelo) laid over UMAP. Individual iCellR clusters scored for a published (**C**) circulating memory, (**D**) resident memory, and (**E**) normalized Cxcr6 expression. (**F**) 15,000 wildtype (WT) and 15,000 *Cxcr6^-/-^* OT-1 T cells were co-transferred into naïve mice and the following day, skin was infected with VACV-OVA. Quantification of the ratio of WT and *Cxcr6^-/-^* OT-1 T cells in spleen and skin and 5 and 14 d.p.i. Data are cumulative from two experiments with n=4. (**G**) Expression of CXCR6 by WT OT-1 T cells in skin relative to spleen over time. Data are representative of at least two experiments with n=3 or 4. (**H**) Experimental design where OT-1 T cells were transferred to naïve mice, which were subsequently infected on the left ear VACV-OVA and the right with VACV (scarification). (**I**) Representative histogram and quantification (**J**) of CXCR6 by OT-1 T cells in spleen or ear skin 14 d.p.i. Geometric mean fluorescence intensity (GMFI). Data are representative of at least two experiments with n=3 or 4. (**K and L**) Nurr77-GFP OT-1 T cells were transferred to naïve mice and the skin was infected with VACV-OVA the next day. (**K**) Representative histograms and (**L**) quantification of CXCR6 by Nurr77-GFP^+^ and Nurr77-GFP^-^ OT-1 T cells in skin 7 d.p.i. Data are representative of two experiments with n=5. (**M**) Expression of CXCR6 by effector CD8^+^ T cells from spleens of listeria monocytogenes (LM-OVA, 7 d.p.i) infected mice restimulated ex vivo with SIINFEKL and TGF-β. Bars represent average +SEM. Data are representative of two experiments with n=3. Cells in skin (**F**) are gated on IV-. Statistical significance determined using paired student’s t test (**F and L**) or one way ANOVA (**G, J and M**), or pairwise Wilcoxon rank test relative to cluster 1 (**C-E**). * p <0.05, ** p < 0.01, *** p < 0.001, **** p < 0.0001.

### Peripheral tissue antigen encounter upregulates CXCR6 expression

The low levels of CXCR6 expression on circulating anti-viral CD8^+^ T cells within the skin (**Fig 1B and 2E**) suggested that CXCR6 expression might not be strictly necessary for infiltration into inflamed skin. To directly test whether CXCR6 is required for tissue infiltration and the early accumulation of anti-viral effector CD8^+^ T cells, we quantified the ratio of co-transferred WT and *Cxcr6*^-/-^ OT-1 T cells at 5 and 14 d.p.i. with VACV-OVA (**Fig 2F**). Importantly, there was no bias toward WT T cells 5 d.p.i. in the draining LN, indicating that CXCR6 deficiency does not impair the magnitude of priming and expansion (**Fig S3A)**. Additionally, both WT and *Cxcr6*^-/-^ OT-1 T cells infiltrated infected skin and were present at a similar ratio to that found in spleen at early (day 5) and late effector (day 14) time points (**Fig 2F**). Consistent with the lack of requirement for tissue entry, CXCR6 expression by WT OT-1 T cells in skin at 5 d.p.i. was almost indistinguishable from expression on WT OT-1 T cells found in the spleen (**Fig 2G**) indicating that CXCR6 was likely upregulated after entry into the skin. Indeed, CXCR6 surface expression increased in skin relative to spleen over time, with established CD8^+^ T_RM_ (day 49) expressing almost 6 times as much surface CXCR6 as their circulating memory counterparts (**Fig 2G**). Consistent with cutaneous VACV infection, CXCR6 was also dispensable for early T cell accumulation in the skin following systemic VSV (**Fig S3B**), and dispensable for accumulation in the small intestine epithelium, and brain 7 d.p.i. WT OT-1 T cells, however, outcompeted *Cxcr6*^-/-^ T cells in the liver, salivary gland, female reproductive tract, and kidney at these early time points (**Fig S3**C), which may indicate either that low levels of CXCR6 on circulating CD8^+^ T cells can support tissue infiltration in a tissue-specific manner, or that CD8^+^ T cells entering these unique tissue environments are more sensitive to loss of CXCR6 expression.

These data indicated that the induction of CXCR6 within skin was a key event for T_RM_ formation. Antigen encounter in peripheral non-lymphoid tissue induces CD69 expression and CD8^+^ T_RM_ formation in both skin (19, 21) and lung (30, 31) following infection. Given the concordance between CXCR6 expression and CD69, we hypothesized that CXCR6 enrichment in skin was also antigen dependent. To test this, we transferred naïve OT-1 CD8^+^ T cells into mice and the next day infected one ear with VACV and the contralateral ear with VACV-OVA. VACV-OVA primed OT-I T cells infiltrate both infected sites independent of antigen expression (19), allowing us to evaluate CXCR6 levels on a fixed CD8^+^ T cell population in the presence or absence of cognate antigen (**Fig 2H**). We observed significantly higher levels of CXCR6 on OT-1 T cells in the presence of cognate antigen (VACV-OVA) compared to VACV infected ears or the spleen 14 d.p.i. (**Fig 2I and J**). Interestingly, while cognate antigen clearly boosted surface CXCR6 expression on anti-viral CD8^+^ T cells, there was a significant increase in CXCR6 on bystander OT-1 T cells in VACV-infected skin relative to spleen, consistent with a role for inflammatory cytokines in surface CXCR6 expression in skin (e.g. IL-2 and IL-15) (24). In order to further confirm a role for antigen encounter and account for potential cytokine differences in VACV and VACV-OVA infected skin, we used mice that report TCR stimulation via transgenic expression of green fluorescent protein (GFP) under control of the *Nr4a1*(Nur77) promoter (Nurr77-GFP) (32). In VACV-OVA infected skin 7 d.p.i., surface CXCR6 was enriched on Nurr77-GFP expressing OT-1 T cells compared to Nurr77-GFP negative OT-1 T cells (**Fig 2K and L),** indicating that recent antigen recognition in skin is associated with increased CXCR6 levels. Furthermore, *ex vivo* activation of CD8^+^ effector T cells (7 d.p.i. listeria monocytogenes-OVA) with cognate SIINFEKL antigen resulted in significant CXCR6 upregulation relative to unstimulated controls over 24 hours (**Fig 2M**). TGF-β signaling is required for the transition of circulating CD8^+^ T cells towards residence, in part by downregulating S1PR1 and upregulating CD103 (33–35), however, the addition of TGF-β did not affect CXCR6 expression in the presence of antigen stimulation *ex vivo* (**Fig 2M)**. These data, together with the observation that CD103^+^ T_RM_ are a subset of CD69^+^CXCR6^+^ T_RM_ (**Fig 1**), indicate that antigen-dependent upregulation of CXCR6 likely precedes TGF-β driven changes and is therefore consistent with CXCR6 as an early feature of CD8^+^ T cell adaptation to the infected dermis.

### CXCR6/ CXCL16 interactions in the dermis do not act to restrain T cell exit

To begin to define the anatomical niche within which CXCR6 acts on CD8^+^ T cell differentiation we analyzed expression of its ligand. CXCL16 was present in skin at all time points analyzed but appeared to peak in expression 7 d.p.i., returning to basal-like levels at late memory time points (**Fig 3A)**. Expression in the skin 7 d.p.i. seemed to largely be driven by myeloid cells, including macrophages and type 1 and 2 conventional dendritic cells (**Fig 3B**). Consistent with the low expression on CD45^-^ cells, at effector time points (14 d.p.i.), CXCL16 was absent from the epidermis (**Fig 3C**), in contrast to epithelial expression observed in the lung (14). Instead, clusters of CXCL16-expressing cells were found within the dermis of infected skin (**Fig 3C**). Given that effector CD8^+^ T cells first enter the dermis through activated capillaries and then transition with time to the epidermal compartment (**Fig 3D**), the enrichment of CXCL16 in the dermis might suggest a role for CXCR6 in the transition to epidermal residence.

**Figure 3.**
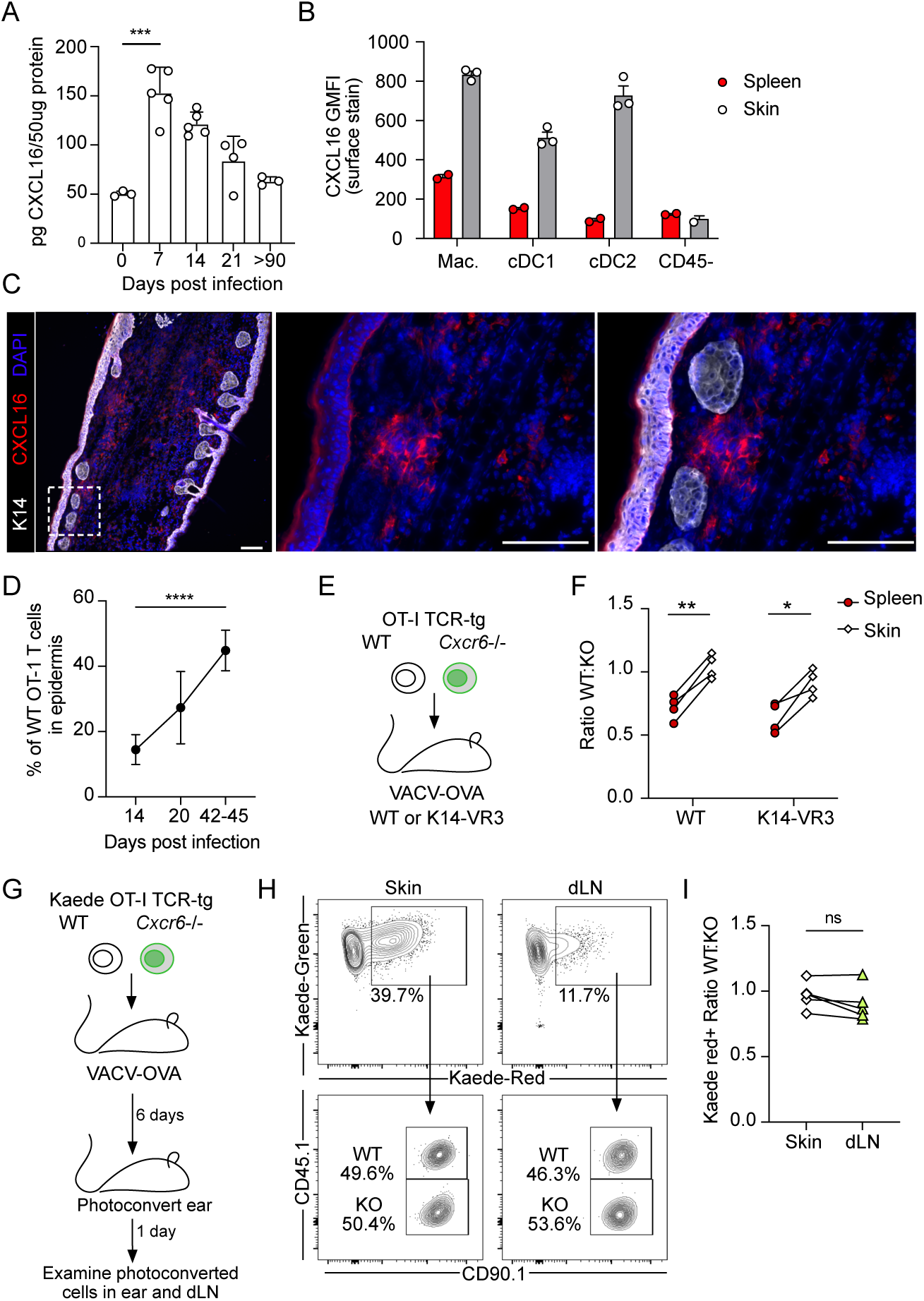
Dermal CXCL16 does not restrain T cell egress out of skin. (**A**) Quantification of CXCL16 in vaccinia virus (VACV) infected skin at various days post infection (d.p.i) by scarification; n=3-5 per group. (B) Quantification of CXCL16 surface expression (GMFI) 12 d.p.i. on dermal and splenic myeloid cells including macrophages (Mac., CD11b^+^F480^+^), conventional dendritic cells type 1 (cDC1, CD11c^+^MHC-II^HI^XCR1^+^) and type 2 (cDC2, CD11c^+^MHC-II^HI^SIRPA^+^). (**C**) Representative immunofluorescence images of skin 14 d.p.i. with VACV. Scale bar = 100μm. (**C**) The percentage of OT-1 T cells in the epidermis after infection with OVA_257-264_-expressing VACV (VACV-OVA) calculated from immunofluorescent images. Data are cumulative of at least two experiments with n=5 (day 14) or n=6 (day 20 and 42-45). (**E**) Experimental design where 15,000 WT and 15,000 *Cxcr6*^-/-^ OT-1 T cells were transferred to K14-VEGFR3-Ig mice and the following day infected with VACV-OVA via skin scarification. (**F**) Quantification of the ratio of WT and *Cxcr6*^-/-^ OT-1 T cells in the spleen and skin of K14 VEGFR3-Ig mice 49 d.p.i. with VACV-OVA. Data are representative from at least two experiments with n=4. (**G**) Experimental design where congenically distinct 15,000 WT Kaede^+^ OT-1 T cells and 15,000 *Cxcr6*^-/-^ Kaede^+^ OT-1 T cells were co-transferred to C57Bl/6J mice and infected the following day with VACV-OVA via skin scarification. Kaede expressing cells in skin were photoconverted 6 d.p.i. and harvested 1 day later from skin and draining lymph node (dLN). (**H**) Representative flow of skin and dLN. (**I**) The ratio of photoconverted WT Kaede red^+^ to photoconverted *Cxcr6*^-/-^ Kaede red^+^ in skin and dLN following photoconversion. Data are representative from at least two experiments with n=5. Statistical significance determined using one-way ANOVA (**A,D**) or paired student’s t test (**F,I**). Error bars represent SD. * p < 0.05, ** p < 0.01, *** p < 0.001, **** p < 0.0001

One hypothesis was that a failure to retain CD8^+^ T cells in the dermis in a CXCL16/CXCR6-dependent manner could lead to increased rates of recirculation and exit through the proximal dermal lymphatic vessels. Dermal lymphatic vessels facilitate CD4^+^ and CD8^+^ T cell egress out of skin (36), and the egress of CD8^+^ T cells limits the number of CD8^+^ T cells that transition to residence (3, 20). We therefore asked if we could rescue *Cxcr6*^-/-^ T_RM_ in the absence of dermal lymphatic vessels, where a physical loss of vessels would prevent T cell exit. Mice in which a VEGFR3-Immunogloubulin fusion protein is expressed under control of the K14 promoter (K14-VEGFR3-Ig), fail to form mature dermal lymphatic vessels (37), restricting both fluid and cellular efflux out of skin (18). K14-VEGFR3 mice exhibit delayed lymphocyte responses to VACV infection but are capable of T cell-mediated viral clearance (18, 20). We again co-transferred equal numbers of congenically distinct, WT and *Cxcr6*^-/-^ OT-1 T cells into naïve K14-VEGFR3-Ig mice and WT controls (**Fig 3E**). At early time points post infection, T cells readily infiltrated the skin of both WT and K14-VEGFR3-Ig mice with no enrichment for WT T cells as seen previously (**Fig. S3D**). At memory time points, however, we expected that if CXCR6 was a retention signal, the removal of the egress route would be sufficient to rescue the accumulation of *Cxcr6*^-/-^ T cells to WT levels in skin. The lack of dermal lymphatic vasculature in K14-VEGFR3-Ig mice, however, failed to normalize the ratio between WT and *Cxcr6*^-/-^ OT-1 T cells in the skin (**Fig 3F**). In line with this finding, *Cxcr6^-/-^* OT-1 T cells were as likely to egress through functional infection-associated lymphatic vessels as their WT counterparts when directly tracked using a photoconvertible transgenic mouse (Kaede-tg) (36, 38). Kaede-tg OT-1 CD8^+^ T cells and Kaede-tg *Cxcr6*^-/-^ OT-1 CD8^+^ T cells were transferred into mice one day prior to infection with VACV-OVA. At 6 d.p.i. the skin was photoconverted and the number of Kaede-red^+^CD8^+^ OT-I T cells were quantified in draining LNs 24 hours later (**Fig 3G, Fig S3E and F**). At this early time point, dermal infiltrating T cells are actively engaging their target and expressing CXCR6 (**Fig. 2G, K and L**). We found no difference in the ratio of photoconverted WT to *Cxcr6*^-/-^ CD8^+^ T cells in the skin compared to those that had egressed to the draining LN (**Fig 3H and I**), and this remained true at later effector time points following transfer into *Rag^-/-^* mice (**Fig. S3G**). These data indicated that despite being a marker of residence, CXCR6 did not appear to be acting as a physical retention cue. Our findings pointed to a dermally-restricted, CXCR6-dependent mechanism of CD8^+^ T cell accumulation in infected skin.

### CXCR6 supports metabolic adaptations during the transition from effector to residence

Our data indicated that CXCR6 likely acted in the dermis to facilitate the transition from effector T cell to T_RM_ over time, though exactly how this chemokine receptor might do this remained unclear. At 21 d.p.i. *Cxcr6* is expressed in all WT OT-1 that bear a transcriptional program consistent with residency (**Fig 2**), indicating its expression is an early event in T_RM_ differentiation. We reasoned that CXCR6 might influence the differentiation potential of effector CD8^+^ T cells in skin. We therefore integrated our WT OT-1 scRNAseq with *Cxcr6*^-/-^ OT-1 T cells extracted from the same skin at 21 d.p.i., when WT and *Cxcr6*^-/-^ OT-1 CD8^+^ T cells are still at roughly equal ratios (**Fig S4A**). Integrated analysis revealed a similar clustering pattern (**Fig 2A**) and psedotime trajectory (**Fig S4B**) as seen for the WT cells alone, again with a circulating cluster (T_CIRC_: cluster 1), 4 clusters of T_RM_ (T_RM_: clusters 2,3,4 and 5) (**Fig 4A**), and a sixth cluster expressing high amounts of *Mcm9* (Mini chromosome maintenance 9) and the transcription factor *Zeb2,* associated with terminal differentiation (39). Interestingly, while both WT and *Cxcr6*^-/-^ OT-1 T cells were present in all clusters the distribution of cells within the T_RM_ clusters was distinct, indicating that the *Cxcr6*^-/-^ T cells were biased in their differentiation trajectory (**Fig 4B**). While the T_CIRC_ cluster 1 scored equally for a published circulating memory score between WT and *Cxcr6*^-/-^, the T_RM_ super cluster demonstrated a reduced capacity to acquire the core T_RM_ transcriptional program in the absence of CXCR6 (**Fig. 4C**) (40). We therefore performed differential gene expression analysis on the T_RM_ supercluster and identified several genes downregulated (e.g. *Icos*, *Il21r*, and *Nr4a3*) (**Fig 4D**) that are functionally implicated in T_RM_ formation within various tissue compartments (40–42). Pathway analysis of the differentially expressed genes revealed that WT cells expressed higher levels of genes involved in leukocyte activation, cytokine signaling and regulation of apoptosis (**Fig 4E**). Genes enriched in these pathways included negative regulators of TCR and NFκB signaling (*Grap, Tnfaip3, Traf1, Nfkbia, Nfkbie*) (43) and components of the immunoproteasome (*Psmb8, Psmb9, Psma6*) important for regulating the transition to memory and cell survival (44). Referencing the TRRUST database confirmed transcriptional regulation of *Nfkb1* (*p* = 8.232e-8) and *Nfkbia* (*p* = 3.476e-5). In contrast, the only pathway significantly upregulated in *Cxcr6*^-/-^ cells was oxidative phosphorylation (**Fig 5E**), largely driven by genes associated with mitochondrial electron transport complex (*Cox6c, Cox7a2, Cox8a, Ndufa4, Ndufb11*). Further analysis of the downregulated genes using HumanBase (45) demonstrated four key modules including NFκB signaling and cellular detoxification, which was driven by reduced expression of *Gstp1*, *Park7*, *Prdx1*, and *Prdx6*. The peroxiredoxin family of antioxidant enzymes reduce hydrogen peroxide and alkyl hydroperoxides and thereby play a protective role in cells. Given that our data predicted that *Cxcr6*^-/-^ T cells exhibited elevated oxidative phosphorylation and an impaired ability to detoxify reactive oxygen species, we hypothesized this could lead to a survival disadvantage within the dermis. Indeed, T cell memory is dependent upon mitochondrial biogenesis, which increases spare respiratory capacity to promote survival and rapid response to recall (46). To validate these transcriptional findings, we directly quantified mitochondrial biomass, membrane potential, and mitochondrial reactive oxygen species in our i*n vivo* co-transfer system. We noted that *Cxcr6*^-/-^ OT-1 T cells exhibited reduced mitochondrial mass (Mito Deep Red, **Fig 4G**), reduced membrane potential (CMX-Ros, **Fig. S4C**), and elevated mitochondrial reactive oxygen species (MitoSoxRed, **Fig. 4H**) 14 d.p.i. relative to WT OT-1 T cells in the same tissue. All consistent with a deficiency in handling of the metabolic byproducts of enhanced oxidative phosphorylation.

**Figure 4.**
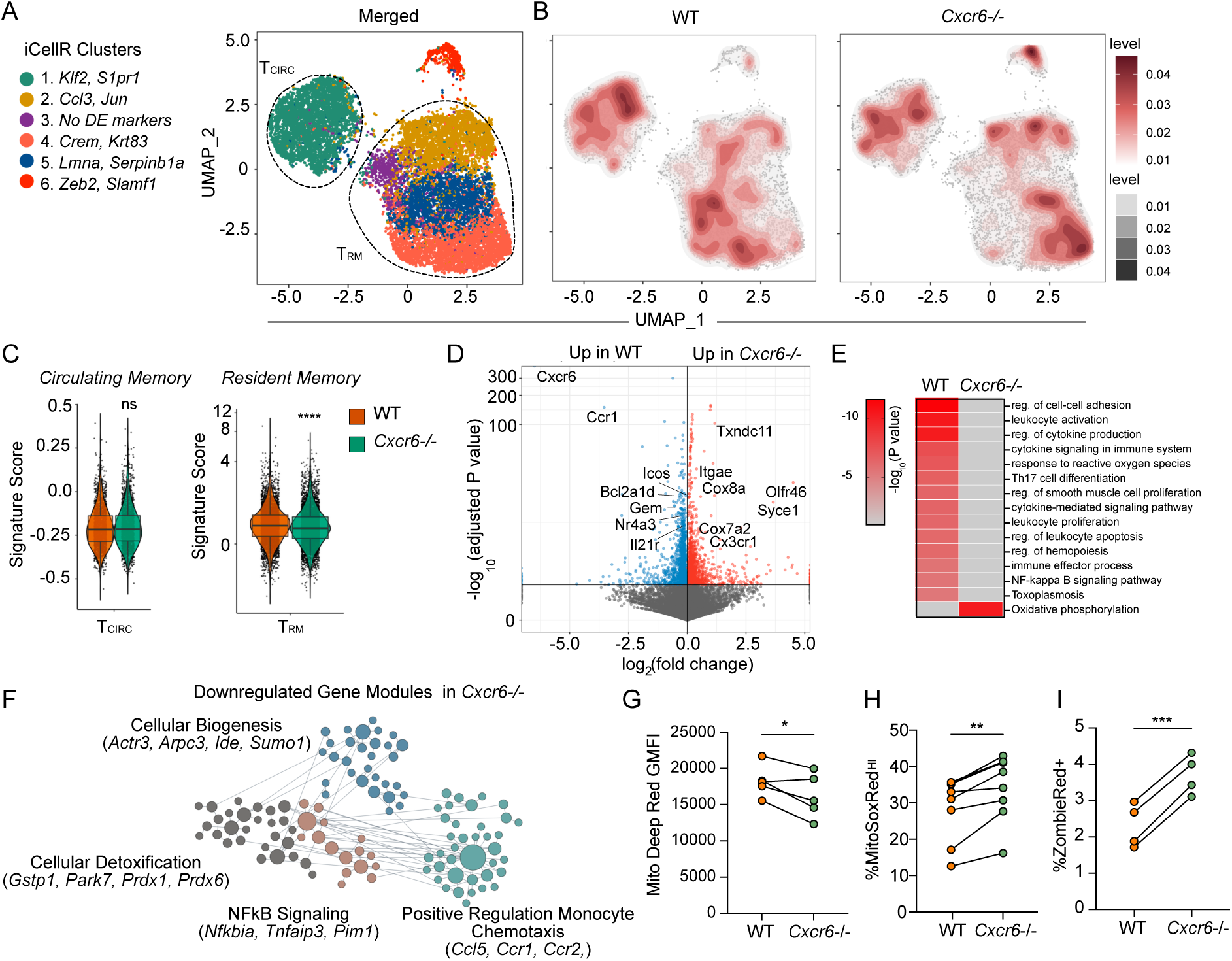
*Cxcr6^-/-^* T cells fail to fully acquire a residence transcriptional program and display signs of metabolic dysfunction and apoptosis. (**A**) UMAP showing merged wildtype (WT) and *Cxcr6*^-/-^ OT-1 T cells isolated from skin 21 days post infection (d.p.i.) with vaccinia virus expressing OVA_257-264_ (VACV-OVA). Resident (T_RM_) and circulating (T_CIRC_) clusters defined as in Fig 2. (**B**) Cell density plots for WT and *Cxcr6*^-/-^ OT-I T cells overlaid on the merged UMAP. (**C**) Cells in T_CIRC_ cluster and T_RM_ clusters scored for published circulating memory (left) or resident memory (right) gene signatures. (**D**) Volcano plot of differentially expressed genes (DEG) between WT and *Cxcr6*^-/-^ cells in combined T_RM_ clusters (2–5). (**E**) Pathway analysis of DEG between WT and *Cxcr6*^-/-^ cells in T_RM_ clusters. p value calculated by hypergeometric distribution test. (**F**) Human Base analysis of gene modules downregulated in *Cxcr6*^-/-^ T_RM_ relative to WT. (**G**) Mito Deep Red GMFI and (**H**) MitoSox Red GMFI in WT and *Cxcr6*^-/-^ OT-I T cells 14 d.p.i. (**I**) Ex vivo uptake of Zombie red viability dye 21 d.p.i. Statistical significance determined using paired student’s t test (**G-I**) or pairwise Wilcoxon rank test (**C**). * p < 0.05, *** p < 0.001, **** p < 0.0001.

**Figure 5.**
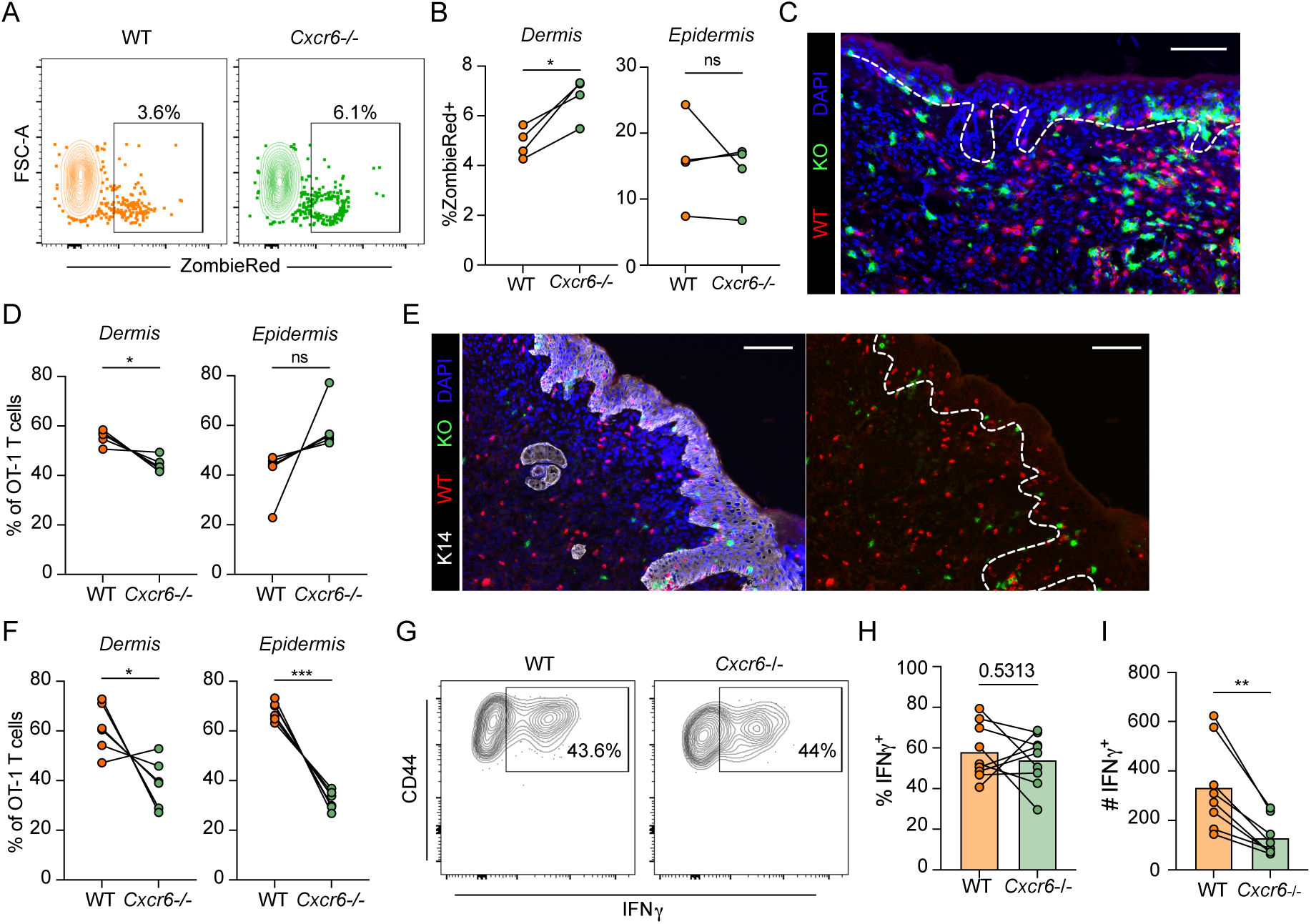
CXCR6 promotes dermal T cell survival during memory formation in infected skin. (**A**) Representative flow plots and (**B**) quantification of ex vivo uptake of ZombieRed viability dye in wildtype (WT) and *Cxcr6*^-/-^ (KO) OT-1 T cells in the dermis and epidermis 14 days post infection (d.p.i) with vaccinia virus expressing OVA_257-264_ (VACV-OVA). Data are representative of at least two experiments with n=4. (**C**) Representative immunofluorescence image and (**D**) quantification of WT and *Cxcr6*^-/-^ (KO) OT-1 T cells in skin at 14 d.p.i. with VACV-OVA. (**E**) Representative immunofluorescence and (**F**) quantification of WT and *Cxcr6*^-/-^ (KO) OT-1 T cells in skin 42-45 d.p.i. with VACV-OVA. White line delineates the epidermis. Scale bar = 100μm. Data are cumulative of two experiments with n=5 (**C,D**) or n=6 (**E,F**). (**G**) Representative flow plots gated on live CD44^+^IV^-^ WT and *Cxcr6*^-/-^ OT-I T cells in VACV-OVA immune mice restimulated with cognate peptide (SIINFEKL; 5 hrs). (**H**) Percent and (**I**) number of IFNγ producing cells from G. Statistical significance determined using paired student’s t test. * p < 0.05, **p<0.01, *** p < 0.001.

Interestingly, in addition to the shifts in transcriptional state, *Cxcr6*^-/-^ T cells were also more represented in Cluster 6 (**Fig 4B**) that exhibited some programs consistent with apoptosis. We reasoned that the dysregulation of reactive oxygen species and mitochondrial fitness could shunt effector T cells off the T_RM_ differentiation pathway resulting in poor survival. Cluster 6 exhibited signs of terminal differentiation, expressing high levels of *Zeb2,* and sat at the end of the predicted pseudotime trajectory defined by RNA velocity (**Fig. S4B**) (47). Following LCMV infection, a *Zeb2*-expressing CD8^+^ T_RM_ cluster did not persist to late memory time points (60-90 d.p.i.) (27), in contrast to a cluster expressing *Id3*, *Jun*, *Fos*, *Ccl3*, and *Ccl4*, genes enriched in our T_RM_ cluster 2, from which cluster 6 arises. Cluster 6 also scored highest for an apoptotic gene signature (R-MMU-109581) (**Fig S4D**), with elevated expression of pro-apoptosis genes such as *Casp8, Bcl211,* and *Acin1* (**Fig S4E**). Even within the T_RM_ clusters, we found that transcripts for the anti-apoptotic proteins, *Bcl2a1b* and *Bcl2a1d,* were significantly reduced in *Cxcr6*^-/-^ relative to WT T cells (**Fig S4F**), perhaps indicating these cells were poised for apoptosis. *Bcl2a1* is an NF-κB target gene with four isoforms that promote lymphocyte survival and differentiation (48). BCL2 protein was indeed lower in *Cxcr6*^-/-^ OT-1 T cells in the skin 21 d.p.i (**Fig 4G and H**), and *Cxcr6*^-/-^ OT-1 T cells exhibited reduced viability in skin relative to matched WT T cells (**Fig 4I**). Taken together, these data indicate that while *Cxcr6*^-/-^ CD8^+^ T cells could adopt a T_RM_-like transcriptional state, they exhibited a deficiency in mitochondrial fitness that may reduce the efficiency with which they transition to long-term residence.

### CXCR6 promotes dermal T cell survival during memory formation in infected skin

The transcriptional data pointed toward a loss of *Cxcr6*^-/-^ T cells as a function of poor survival likely driven by a combination of effects including reduced cytokine signaling, metabolic dysfunction, and a failure to fully activate a resident memory transcriptional program. Given the high expression of CXCL16 in the dermis, we asked whether CXCR6 preferentially impacted dermal CD8^+^ T cell viability and thereby transition to long-term maintenance in the epidermis. At 14 d.p.i. with VACV-OVA, when OT-1 T cells are present in both the dermis and epidermis, *Cxcr6*^-/-^ T cells exhibited similar viability in the epidermis but reduced viability in the dermis relative to WT (**Fig 5A and B**). We next asked if the reduced viability of *Cxcr6*^-/-^ T cells in the dermis impacted their frequency in the dermis and epidermis over time. Using immunofluorescent imaging of skin 14 d.p.i., we observed that WT OT-1 T cells were more prevalent than *Cxcr6*^-/-^ OT-1 T cells in the dermis, while in the epidermis, WT and *Cxcr6*^-/-^ OT-1 T cells were present at roughly equal levels (**Fig 5C and D**). This suggested that WT cells were outcompeting *Cxcr6*^-/-^ in the dermis at effector time points, prior to acquiring epidermal residence. We therefore hypothesized that CXCR6 supported tissue adaptation and survival during the transition to residency. Consistent with this hypothesis, at later time points, 42-45 d.p.i, *Cxcr6*^-/-^ OT-1 T cells were now less prevalent in both the dermis and epidermis (**Fig 5F**). These data indicate that *Cxcr6*^-/-^ CD8^+^ T cells form T_RM_ at suboptimal levels due to reduced survival in the dermis during a critical window of T_RM_ differentiation. Finally, we asked whether once established, if loss of CXCR6 limited the functionality of T_RM_. Notably, even at even very late time points, > 100 d.p.i, the ratio between WT and *Cxcr6*^-/-^ T cells remained approximately 2 to 1 (**Fig. 1G**), indicating that CXCR6 might be dispensable for long-term maintenance. Furthermore, restimulation of established T_RM_ with cognate peptide induced equal amounts of IFNγ per cell in both WT and *Cxcr6*^-/-^ OT-I T cells (**Fig. 5G and H**), further supporting the interpretation that CXCR6 promotes the efficiency of T_RM_ formation but is not required for their long-term maintenance or function. Our data therefore positions CXCR6 as a TCR-dependent, interstitial signal necessary for optimal survival during the transition to residence but dispensable for long-term maintenance and effector function.

## DISCUSSION

The formation of T_RM_ depends on mechanisms of tissue adaptation, cellular positioning, and differentiation. While the transcriptional programs that are enriched during transition from effector to memory are well described, the functional implications of key markers remain unclear. Transcriptional analyses of CD8^+^ T cells in diverse anatomical locations of mice and humans identified the chemokine receptor CXCR6 as a highly expressed marker of T_RM_ (5–7). In this study, we investigated the functional significance of CXCR6 for T_RM_ formation. Using competitive co-transfer experiments of *Cxcr6*^-/-^ and WT controls, we find that CXCR6 expression is needed for optimal T_RM_ formation in peripheral non-lymphoid tissues after viral infection. Surprisingly, we found that CXCR6 has no effect on expansion in the draining lymph node, early accumulation of CD8^+^ T cells in the skin, nor does it prevent egress out of skin after infection. Instead, CXCR6 deficiency diminished T_RM_ quantity by reducing CD8^+^ T cell viability in inflamed skin, specifically within the dermis, leading to a failure to transition to epidermal residence.

Using adoptive transfer experiments and multiple viral models we show that CXCR6-deficiency diminished the expected quantity of T_RM_ in diverse tissues including brain, kidney and skin. While chemokines and their receptors are known to play critical roles in shaping the spatial distribution of T cells between and within tissues, here we identify a migration-independent mechanism of action. Across various non-lymphoid peripheral tissues and two infections, CXCR6 was not absolutely required for tissue infiltration. This is similar to observations made in mouse models of influenza infection, where CXCR6 is not needed for T cell entry into the lung parenchyma (14), and HSV infection of skin, where *Cxcr6*^-/-^ CD8^+^ T cells still form suboptimal T_RM_ numbers even after direct intradermal injection (49). Furthermore, CXCR6 did not appear to help anchor T cells in tissue, as deficient T cells did not exhibit increased rates of lymphatic egress and loss of the lymphatic vasculature was not sufficient to boost T_RM_ formation. Further, we find CXCL16 to be most highly expressed in the dermis at effector time points after infection, which pointed towards a functional contribution prior to acquisition of epidermal residence. This observation was interestingly in contrast to the lung where CXCL16 is expressed by the airway epithelium and thought to promote antigen-specific CD8^+^ T cell homing to that tissue compartment (14). Interestingly, in VACV-infected skin, CXCR3-deficient T cells fail to properly migrate towards and kill infected cells in the epidermis (50), which may hint at tissue and or viral-specific mechanisms of target cell homing.

Our findings therefore point to a model where CD8^+^ T cell antigen encounter in the dermis and concomitant CXCR6 upregulation precedes migration to the epidermis. Surface expression of CXCR6 by CD8^+^ T cells in skin is upregulated between 5 and 7 d.p.i. and depends on local antigen recognition. This is consistent with recent work that demonstrates that TCR signaling strength regulates *Blimp1* expression, which thereby tunes the chemotactic features of effector CD8^+^ T cells during the transition to residence (21). Similarly, we recently demonstrated in tumors that local antigen encounter downregulates CXCR4, which acts as a mechanism of tumor egress (29), and in concordance *Cxcr4* was transcriptionally downregulated in cells transitioning to residence following infection. In contrast, and as expected based on data presented here, CXCR6 was enriched on tumor-retained CD8^+^ T cells that exhibited signs of chronic antigen stimulation (22, 29). Interestingly, tissue entry and initial antigen presentation may be tightly linked in infected skin presenting the possibility that *Cxcr6* upregulation occurs rapidly in a dermal, perivascular niche (51), where at least in tumors, the CXCL16-expressing dendritic cells reside (10). We found that *Cxcr6* was expressed across transcriptional states in resident memory precursors and followed a similar kinetic as antigen-dependent induction of CD69 (19). Further consistent with our model, CXCR6 expression on restimulated effector CD8^+^ T cells was not affected by the presence of TGF-β and CXCR6 upregulation precedes TGF-β-dependent CD103 upregulation by several days. CD103 permits adhesion to E-cadherin expressed in epidermal keratinocytes and TGF-β is activated at the epidermal interface (52, 53). While TGF-β may facilitate residence in part through its role in downregulating *S1pr1, S1pr5* and *Cxcr4* (35), our data suggests CXCR6 function precedes TGF-β dependent mechanisms of residency. Taken together, these findings indicate that CXCR6 upregulation likely takes place in the dermis, early after tissue entry and before CD8^+^ T cell transition to the epidermis.

Our data reveal that CXCR6 plays an important role for CD8^+^ T cells in the transition from circulating effector to resident memory but CXCR6 deficiency does not place an absolute block on T_RM_ differentiation. *Cxcr6*^-/-^ OT-I T cells at an early memory time points adopt the same transcriptional states seen in WT OT-1 T cells taken from skin, we did find that CXCR6-deficient T_RM_ in the skin score lower for a resident memory gene signature. The differences in this resident memory gene signature, while modest, were driven by reduced expression for over a dozen genes including several genes with known roles in T_RM_ formation including, *Il21r, Nr4a3* and *Icos* (40–42). Decreased expression of *Il21r* is particularly interesting as IL-21 promotes memory T cell survival and proliferation, in a similar manner to IL-15 (54). In addition to these known T_RM_-related transcripts, *Cxcr6*^-/-^ T cells appear to exhibit dysregulation of NFκ-B signaling, the immunoproteasome, oxidative phosphorylation and apoptosis, all of which may indicate a cell not activating the appropriate cues to efficiently transition to memory. Consistent with the elevations in oxidative phosphorylation but reduced antioxidant capacity, *Cxcr6-/-* T cells accumulated more mitochondrial reactive oxygen species when compared to WT OT-I T cells from the same infected skin. It seems likely that these observed changes converge on multiple survival and activation pathways that decrease the probability of T cell survival and therefore long-term residence. These observations altogether may indicate that CXCR6 functions to retain activating CD8+ T cells in transient niches that optimize their receipt of survival cues to increase the efficiency of T_RM_ formation.

Previous studies evaluating the importance of CXCR6 expression by CD8^+^ T cells in anti-tumor responses shed further light on why CXCR6 is crucial for T_RM_ development. In mouse models of melanoma and pancreatic cancer, CXCR6 expression is necessary for tumor control by CD8^+^ T cells (10, 23). In these systems, CXCR6 optimizes interactions with CXCL16 and IL-15 expressing DCs in perivascular niches of the tumor stroma (10). T_RM_ dependency on IL-15 varies by anatomical location (55) but IL-15 is essential for the maintenance of CD103^+^ T_RM_ in skin (2). IL-15 regulates oxidative phosphorylation for CD8^+^ T cells by inducing mitochondrial biogenesis and also increases BCL-2 expression (46). Our data associated loss of BCL-2 with a reduction in mitochondrial biogenesis and membrane potential, which could lead to the increased production of reactive oxygen species that we see and ultimately cell death. Consistently we see early cell death in the dermis but not the epidermis, indicating that balancing these signals first in the dermis is crucial for long-term epidermal outcome. These data are consistent with a hypothesis that reduced exposure to IL-15 in the dermis impairs the metabolic reprogramming necessary for survival and transition to memory. We cannot, however, exclude the possibility that CXCR6 might directly activate NF-κB through Akt (56), thereby improving CD8^+^ memory T cell fitness in part through direct regulation of BCL2 (57).

Interestingly, in other peripheral tissues, such as the kidney, where T cells experience elevated levels of hypoxia (58), these stress-adaptation mechanisms may have more significant effects on interstitial persistence. While *Cxcr6*^-/-^ T cells were equally competent to reside and accumulate in skin at early time points (7 d.p.i.), they were already significantly impaired in the kidney, salivary gland, liver, and female reproductive tract at this same time point. While additional work would be needed to rule out a direct effect of CXCR6 on tissue homing and transendothelial migration, the relatively low levels of expression on circulating CD8^+^ T cells continue to support the hypothesis that CXCR6 helps meet the demands of the interstitial tissue microenvironment. The highly conserved expression of CXCR6 on resident, NK, and unconventional T cells (5, 6, 59–61), all suggests it likely facilitates a critical mechanism of cellular adaptation adopted across peripheral non-lymphoid tissues and inflammatory states.

Taken all together, our findings demonstrate that CXCR6 instigates mechanisms of cellular adaptation to tissue and promotes local survival by facilitating signals that boost mitochondrial biogenesis and protect cells from death. Here, we have demonstrated that CXCR6 contributes to the process of T_RM_ formation by facilitating CD8^+^ T cell survival and transition to residency during phases of immune resolution 14-30 days post infection. It will be interesting to determine if CXCR6 remains required to maintain T_RM_ survival in the epidermis or if it is only needed during these inflammatory periods when CXCL16 expression is elevated, dermal effector cells are transitioning to epidermal memory, and survival cytokines might be limiting. Experimental models allowing for spatial and temporal control of CXCR6 or its ligand, CXCL16, will be needed to determine their role in maintenance and potential to regulate established T_RM_ populations. Since T_RM_ are potent sources of localized immunity that can mediate tissue-specific autoimmune responses, mechanisms of cellular adaptation to tissue stressors that are required for T_RM_ formation and maintenance may represent targets to limit T cell-induced injury across tissue sites (58). In sum, our data provide mechanistic insight into the parameters that tune CD8^+^ T cell transition to residence and contributes to an emerging hypothesis that targeting T cell position is a viable strategy for tissue-specific immunotherapy across disease indications.

## MATERIALS AND METHODS

### Mice

C57BL/6J, B6.SJL-PtprcaPepcb/BoyJ (CD45.1), C57BL/6-Tg(TcraTcrb)1100Mjb/J (OT-1), B6.129P2-Cxcr6tm1Litt/J (*Cxcr6*^-/-^) (24), B6.PL-Thy1a/CyJ (CD90.1), C57BL/6-Tg(Nr4a1-EGFP/cre)820Khog/J (Nur77-GFP) mice were purchased from Jackson Laboratory and bred in specific pathogen-free conditions at Oregon Health and Sciences University or New York University. B6.Cg-Tg(CAG-tdKaede)-15Utr (Kaede-Tg) (62) were obtained via D.J. Fowell in agreement with RIKEN BioResource Research Center. K14-VEGFR3-Ig mice (37) were a generous gift from Dr. Kari Alitalo and obtained from Dr. Melody A. Swartz. Mice were age and sex matched and female mice were used unless otherwise stated. OT-1 chimeras were created by adoptively transferring 15,000 CD8^+^ T cells from OT-1 mice into naïve mice. For co-transfer experiments, 15,000 wildtype OT-1 T cells and 15,000 *Cxcr6*^-/-^ OT-1 T cells were co-transferred into naïve mice. For experiments with K14-VEGFR3-Ig mice, both male and female recipients were used and 30,000 wildtype OT-1 T cells and 30,000 *Cxcr6*^-/-^ OT-1 T cells were co-transferred into naïve mice. Mice were used between 8-14 weeks of age. All animal procedures were approved and performed in accordance with the Institutional Animal Care and Use Committees at OHSU and NYU Langone Health.

### Pathogens and Infections

Mice were infected the following day after adoptive transfer, with vaccinia virus (VACV), VACV expressing SIINFEKL (VACV-OVA) or VSV expressing full length ovalbumin (VSV-OVA). VACV, and VACV-OVA were propagated in BSC-40 cells as per standard protocols. Mice were infected by administering 1-5 x10^6^ PFU of VACV, or VACV-OVA in 10 μl of PBS to the ventral side of the ear pinna followed by 25 pokes with a 29-G needle (skin scarification). For VSV-OVA infections, mice were injected intravenously with 1 x 10^7^ PFU. For reactivation of T_RM_ with peptide, VV-OVA immune mice were injected RO with 300 μl of 0.5mg/ml BFA in PBS. 30 minutes later, cells in blood were labeled with RO injection of 3 μg of anti-CD45 APC. Immediately after IV labeling, 25 μg of SIINFEKL in 10ul of 4:1 Acetone:DMSO was placed on both sides of ear skin and poked 10-15 times with a needle. Mice were taken down 5 hours later. To generate effector CD8^+^ T cells for in vitro studies, ActA deficient *Listeria monocytogenes* expressing SIINFEKL (LM-OVA) was grown in tryptic soy broth supplemented with 50 μg/ml streptomycin at 37°C and until it reached 1 x 10^8^ CFU/ml (OD600=0.1). 1 x 10^7^ CFU was administered intravenously and spleens harvested 7 days later.

### In vitro Restimulation

Splenocytes were cultured *in vitro* for 24 hours in 1nm SIINFEKL peptide or with 10ng/mL of TGF**-**β in RPMI containing 10% FBS.

### In vivo T cell egress

To label Kaede expressing T cells, ear skin was photoconverted 6 days after infection using 405-nm light (Steele 2019) for 2 minutes (1 minute each side) at 10mW. Mice were euthanized 24 hours later.

### Immunofluorescent microscopy

After euthanizing mice, ears were frozen in OCT on an isopentane bath. 7μm sections were cut and fixed in acetone for 15 minutes. Slides were blocked for 10 minutes with (1% BSA in PBS) and stained with antibodies in 1%BSA in PBS for 1-2 hours at room temperature. Antibodies were obtained from Biolegend, Invitrogen, BD Biosciences, Bioss, and Tonbo and included CXCL16 (pAb Bioss), K14 (POLY9060), CD8 (53-6.7), CD45.1 (A20), CD90.1 (OX-7). Nuclei were detected and cover slips were attached using 4′,6-diamidino-2-phenylindole dihydrochloride (DAPI) containing Prolong Diamond Anti-Fade (Life Technologies). Slides were imaged on a Keyence BX-X810 microscope.

### Leukocyte isolation

Ears of mice were removed and the dorsal and ventral sides separated and incubated for 45 minutes at 37 C in 1ml of HBSS (Hyclone) containing CaCl_2_ and MgCl_2_ supplemented with 125 U/ml of collagenase D (Invitrogen) and 60 U/ml of Dnase-I (Sigma-Aldrich). Tissue was smashed on a scored plate and poured through a 70 μm filter. For some experiments, leukocytes were purified by resuspending pellets in 35% Percoll (GE Healthcare) and HBSS followed by room temperature centrifugation at 1800 RPM. Single cell suspensions of leukocytes from lymph nodes and spleens were made by smashing the tissue through a 70μm filter. Spleens were resuspended for 2 minutes in 2ml of ammonium-chloride-potassium lysis buffer. Lymphocyte isolation from all other non-lymphoid tissues was performed as previously described (63)(Steinert 2015) using 35% Percoll and HBSS followed by room temperature centrifugation at 1800 RPM.

### Flow Cytometry

For intravenous (IV) labeling, mice were injected intravenous with 2-3 μg of anti-CD45 antibody and euthanized 3 minutes later. For surface stains, single-cell suspensions were stained with antibodies in FACS buffer (1%BSA in PBS) for 30 minutes at 4C. Antibodies were obtained from Biolegend, Invitrogen, BD Biosciences, BD Pharmingen, Bioss, and Tonbo and included CXCL16 (12–81), BCL2 (BCL/10C4 or 3F11), CD8 (53-6.7), CD45.1 (A20), CD90.1 (OX-7), CD90.2 (30-H12), CXCR6 (SA051D1), CD69 (H1.2F3), CD103 (2E7), CD44 (IM7), CD45 (30-F11), and CD16/32 (2.4G2). Ghost dye 780 (ZombieRed) was used to measure cell viability. For some flow cytometry analysis, *Cxcr6*^-/-^ CD90.1^+^ OT-1 T cells were identified via GFP expression and AF488-CD90.1 antibody. Mitochondrial staining was performed as per manufacturer (Invitrogen) instructions. Staining of intracellular BCL2 was performed with True-Nuclear™ Transcription Factor Buffer Set (Biolegend) as per manufacturer protocol. Samples were analyzed on a BD LSR II, or BD FACSymphony. FlowJo 10.8.1. was used to analyze flow cytometry data.

### ELISA

Ears from naïve or previously infected mice were weighed and snap frozen in liquid nitrogen. Ears were homogenized and CXCL16 concentration was quantified using Mouse CXCL16 DuoSet ELISA (R&D Systems) as per the manufacturers protocol.

### Sorting and single-cell RNA-seq

Ears of C57Bl/6J mice were infected with VACV-OVA the day after receiving 15,000 WT OT-1 T cells and 15,000 *Cxcr6*^-/-^ OT-1 T cells. At 21 days post infection, lymphocytes were isolated from skin after intravenous injection of anti-CD45-PE antibody as previously described. WT OT-1 T cells (CD8^+^CD90.2^+^CD44^+^CD45 IV^-^CD45.1^+^CD90.1^-^) and *Cxcr6*^-/-^ OT-1 T cells (CD8^+^CD90.2^+^CD44^+^CD45 IV^-^CD45.1^-^CD90.1^+^) were sorted. The sorted cellular suspensions were loaded on a 10x Genomics Chromium instrument to generate single-cell gel beads in emulsion. Libraries were prepared using Single cell 3’Reagent kits v3.1 (Chromium Next GEM Single Cell 3’ GEM, Library & Gel Bead Kit v3.1, 16 rxns PN-1000121;10x Genomics) and were sequenced using Illumina Novaseq 6000.

### Bioinformatics and pathway analysis

Quality control was performed on sequenced cells to calculate the number of genes, UMIs and the proportion of mitochondrial genes for each cell. Cells with low number of covered genes (gene-count < 200) and high mitochondrial counts (mt-genes > 0.08) were filtered out. There were 8634 cells in the *Cxcr6*^-/-^ sample and 7409 cells in the WT sample after filtering. The matrix was normalized based on their library sizes. A general statistical test was performed to calculate gene dispersion, base mean and cell coverage to use to build a gene model for performing Principal Component Analysis (PCA). Genes with high coverage (top 500) and high dispersion (dispersion > 1.5) were chosen to perform PCA and batch alignment using iCellR R package (v1.6.5). T-distributed Stochastic Neighbor Embedding (t-SNE) and Uniform Manifold Approximation and Projection (UMAP) were performed on the top 10 PCs. PhenoGraph (64) clustering was then performed on the UMAP results. Marker genes were found for each cluster and visualized on heatmaps, bar plots and box plots. The marker genes were used to determine cell types. Proportion (percentage) of the cell communities in each condition were calculated. The Tirosh scoring method (65) was used to calculate gene signature scores for circulating and resident memory (40). RNA velocity results were performed by using velocyto (v0.17) (66) https://doi.org/10.1038/s41586-018-0414-6) and scVelo (v0.2.3) (https://scvelo.readthedocs.io/). Pathway analysis was performed using marker genes in Metascape v.3.5 (http://metascape.org).

### Statistics

Statistical analysis was performed using GraphPad Prism 9 software. Comparisons between two groups were conducted using paired or unpaired students t tests. Comparisons between three or more unpaired groups were conducted using ANOVA with Tukeys multiple comparison test. Comparisons between three or more paired groups were conducted using repeated measures ANOVA with Greenhouse-Geisser correction. Proper sample size was based off of prior experience. Statistical analysis of sequencing data was performed with pairwise Wilcoxon rank test and Bonferroni correction in R. P values are reported as follows: * p < 0.05, ** p < 0.01, *** p < 0.001, **** p < 0.0001. Error bars show the mean ±SD.

## Supplementary Materials

Figs. S1 to S4

Table S1 to S2

## Acknowledgments

The authors would like to thank Niroshana Anandasabapathy and Susan R. Schwab for critical feedback on the manuscript, and acknowledge all members of the Lund Lab for critical feedback and technical support.

## Funding

National Institutes of Health grant R01CA238163 (AWL)

National Institutes of Health grant T32AI100853 (TAH)

National Institutes of Health grant T32CA106195 (MMS)

Cancer Research Institute, Lloyd J. Old STAR Award (AWL)

American Cancer Society, RSG-18-169-01-LIB (AWL)

National Institutes of Health grant P30-CA016087 (Laura and Isaac Perlmutter Cancer Center supporting the Flow Cytometry and Cell Sorting Core and the Genome Technology Center (RRID: SCR_017929)

National Institutes of Health grant P30CA069533 (OHSU Knight Cancer Center supporting the Flow Cytometry and Cell Sorting Core)

## Author contributions

Conceptualization: TAH, AWL

Methodology: TAH, ZL

Investigation: TAH, OI, ZL, ACS, MMS, TM

Funding acquisition: AWL, TAH

Supervision: AWL

Writing – original draft: TAH, AWL

Writing – review & editing: TAH, OI, ZL, ACS, MMS, TM, AWL

## Data and materials availability

scRNAseq data is deposited in GEO, GSE223727. All sequence data, code, and materials used in the analysis will be made publicly available upon manuscript acceptance.

