## Supplemental Figures for "CXCR6 promotes dermal CD8^+^ T cell survival and transition to long-term tissue residence"

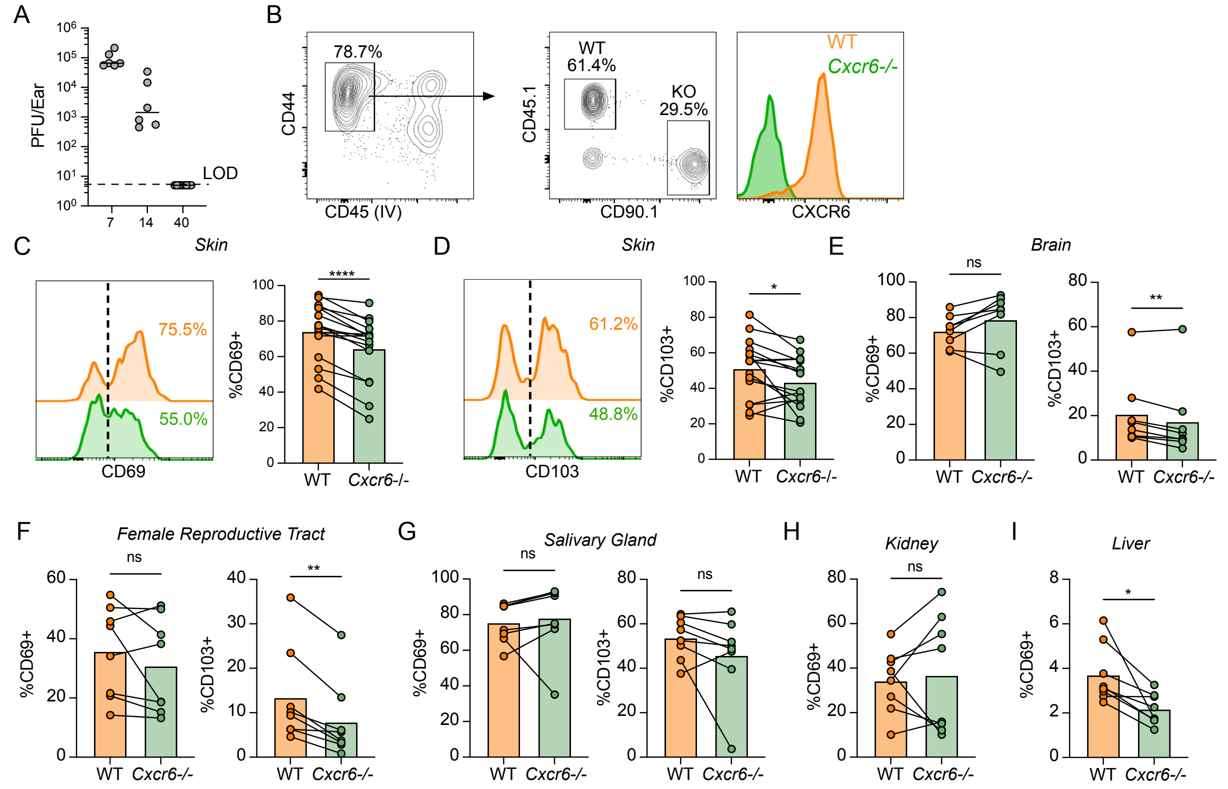


**Supplementary Fig 1. CXCR6 boosts the T$_{RM}$ fate across peripheral, non-lymphoid tissues. (A)** Viral titers (plaque-forming units, PFU) following vaccinia virus (VACV) infection by scarification. Limit of detection = LOD. (**B**) Representative gating strategy used to identify WT OT-1 T cells and *Cxcr6*$^{-/-}$ OT-1 T cells from non-lymphoid tissues (brain after VSV-OVA is shown). (**C** and **D**) Representative flow analysis and quantification of CD69 (**C**) and CD103 (**D**) on OT-I T cells in skin at least 49 days post infection (d.p.i) with vaccinia virus (VACV). Data are cumulative from 3 experiments for a total of n=12. (**E-G**) CD69 (left) and CD103 (right) expression on OT-1 T cells in brain (BRN, **E**), female reproductive tract (FRT, **F**), and salivary gland (SG, **G**) 31-33 d.p.i with VSV-OVA. (**H and I**) CD69 expression on OT-1 T cells in kidney (KD, **H**) and liver (LVR, **I**) 31-33 d.p.i with VSV-OVA. Statistical significance was determined using paired student's t test (**C-I**). * p <0.05, ** p <0.01,

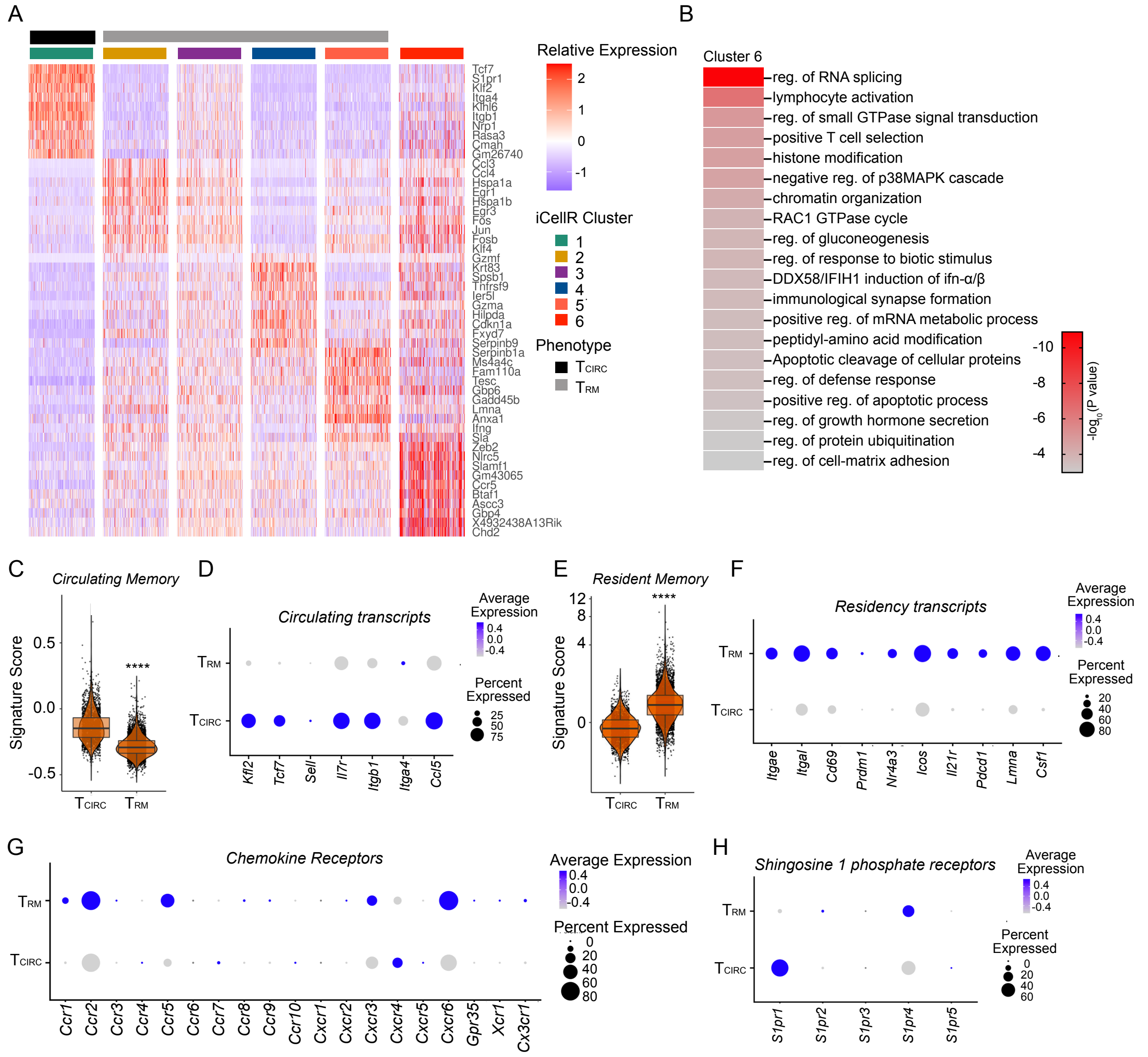


**Supplementary Fig 2. scRNAseq of circulating and resident memory T cells following vaccinia infection in skin.** (**A**) Imputed heatmap of differentially expressed genes from OT-1 T cell scRNA-seq 21 days post infection (d.p.i.). (**B**) Pathway analysis of differentially expressed genes in cluster 6. P value calculated by hypergeometric distribution test. (**C**) Scoring of circulating memory ($T_{CIRC}$) published gene signature and (**D**) normalized expression of select $T_{CIRC}$ transcripts. (**E**) Scoring of resident T cell memory ($T_{RM}$) published gene signature and (**F**) normalized expression of select $T_{RM}$ transcripts. $T_{CIRC}$, cluster 1 and $T_{RM}$, clusters 2-5.Normalized expression of chemokine receptors (**G**) and sphingosine phosphate receptors in $T_{RM}$ and $T_{CIRC}$. (**C and E**) statistical significance determined using pairwise Wilcoxon rank test (**C and E**). *** $p < 0.001$.

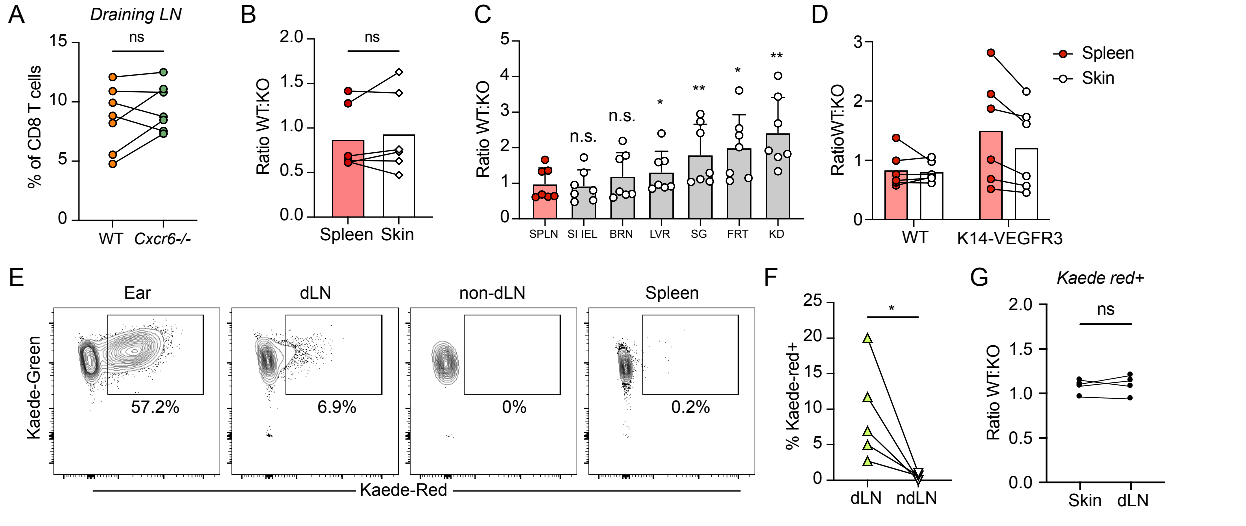


**Supplementary Fig 3. CXCR6 expression is not required for priming, early accumulation, or egress form skin.** (**A**) Prevalence of WT and *Cxcr6*$^{-/-}$ OT-1 T cells in the draining lymph nodes (LN) 5 days post infection (d.p.i.) with VACV-OVA via scarification. (**B**-**C**) Ratio of WT and *Cxcr6*$^{-/-}$ OT-1 T cells in spleen and skin 7-8 (skin, **B**) or 7 d.p.i. (peripheral non-lymphoid tissues; **C**) with VSV-OVA (Spleen, SPLN; small intestine interepithelial lymphocytes, SI IEL). Gated on IV$^{-}$ in all tissues except spleen and liver. (**D**) Ratio of WT and *Cxcr6*$^{-/-}$ OT-I T cells 14-15 days post VACV infection in K14-VEGFR3-Ig mice or littermate, wildtype controls (WT). Cumulative data from two experiments. (**E**) Representative flow plots and quantification (**F**) from experimental design in Fig. **3G** showing photoconversion of Kaede$^{+}$ WT and *Cxcr6*$^{-/-}$ OT-1 T cells in tissue. Gated on CD8$^{+}$CD44$^{+}$CD90.1$^{+}$. Draining (cervical; dLN) and non-draining lymph node (contralateral cervical; ndLN). (**G**) WT and *Cxcr6*$^{-/-}$ Kaede$^{+}$ OT-I T cell egress from infected skin in Rag$^{-/-}$ mice. Photoconverted 13 d.p.i and analyzed 14 d.p.i. Data are cumulative of two experiments with n=7 (**B and C**), n=6 (**A**), n=5 (**F**), or n=4 (**G**). Statistical significance was determined using paired student's t test (**A, B, D, F and G**) or repeated measures one-way ANOVA (**C**). * $p < 0.05$, **$p<0.01$.

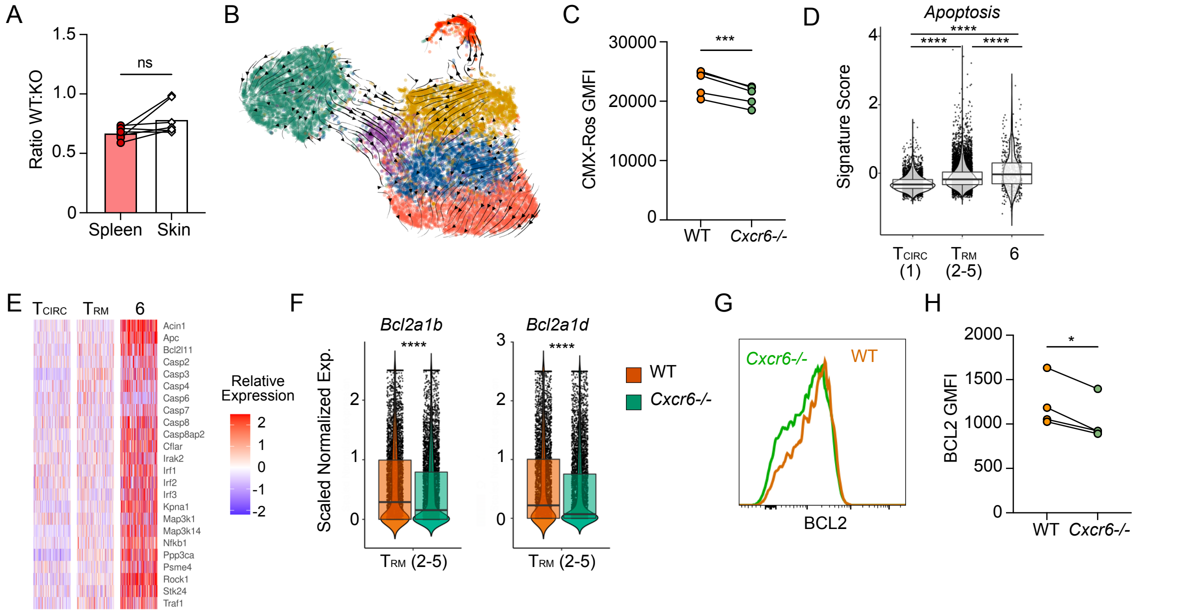


**Supplementary Fig 4. Transcriptional and phenotypic differences in CXCR6$^{-/-}$ OT-1 T cells.** (**A**) Ratio of WT and *Cxcr6*$^{-/-}$ OT-1 T cells 21 days post infection (d.p.i.) with vaccinia virus expressing the immunodominant CD8$^{+}$ epitope in ovalbumin (VACV-OVA) via scarification. Data are cumulative from two experiments with n=7. (**B**) Projected RNA velocity (scVelo) laid over UMAP. (**C**) GMFI of CMX-Ros on OT-1 T cells in skin 14 d.p.i. with VACV-OVA. Data are representative of two experiments with n=4 for each experiment. (**D**) Integrated (WT and *Cxcr6*$^{-/-}$) clusters were scored for apoptosis signature (R-MMU-109581). (**E**) Expression of select apoptosis-related transcripts in integrated clusters (Bioplanet 2019). (**F**) Scaled and normalized expression of *Bcl2a1* isoform transcripts by cells in T$_{RM}$ clusters. (**G and H**) WT and *Cxcr6*$^{-/-}$ OT-1 T cells were isolated from VACV-OVA infected ear skin 21 d.p.i. (**G**) Representative flow and (**H**) quantification of BCL2 GMFI. Statistical significance was determined using paired student's t test (**A,C, and H**)), one-way ANOVA (**D**), or pairwise Wilcoxon rank test (**F**). *p<0.01, *** p < 0.001, **** p < 0.0001.
